# Link Your Sites (LYS.py): Coupling your PAML codeml results and homologous protein structures in PyMOL

**DOI:** 10.1101/380394

**Authors:** Lys Sanz Moreta, Rute Andreia Rodrigues da Fonseca

**Affiliations:** København Universitet. Bioinformatics Center; Natural History Museum of Denmark- Statens naturhistoriske museum

## Abstract

The visualization of the molecular context of an amino acid mutation in a protein structure is crucial for the assessment of its functional impact and to understand its evolutionary implications. Currently, searches for fast evolving amino acid positions using codon substitution models like those implemented in PAML (Z. Yang, 2000) are done in almost complete proteomes, generating large numbers of candidate proteins that require individual structural analyses. Here I present a python wrapper script that integrates the output of PAML with the PyMOL visualization tool to automate the generation of protein structure models where positively selected sites are mapped along with the location of putative functional domains.

## INTRODUCTION

Studies in genetic evolution infer positive selection occurring at specific codons (amino acids) in the DNA (protein) sequence thanks to models, such as PAML site models (Z. Yang, 2000), that evaluate positive selection occurring at certain sites. PAML branch-site models (J. Zhang, 2005) can also identify clades or branches in the tree that are evolving under stronger positive selection conditions than the rest of the tree.

Positive selection is evaluated through the ω value that calculates the ratio between the amount of non-synonymous mutations per non-synonymous site and the amount of synonymous mutations per synonymous site. Non-synonymous mutations can be relevant if the amino acid switch introduced generates a change in the physico-chemical properties of the residue and consequently affects the protein 3D structure. An approach to evaluate the impact of these mutations in the structure is to observe them in homologous protein structures and analyze if they are located within the functional domain residues. Mutations in the functional domain are more likely to affect the protein’s function than if they are located in other parts of the structure.

Proteins structures are highly conserved due to their direct association with the protein’s function. Protein structure is defined by the amino acid chain derived from the DNA sequence transcription and translation. The amino acids form a backbone that is folded into a specific conformation regarding its interactions with the external medium, mainly aqueous, in which is submerged. The folding patterns are dictated by a series of non-covalent bonds *(hydrogen bonds, ionic bonds* and *van der Waals attractions)* directed by the residue’s side chains. If the residues in the functional domain are exchanged with an amino acid with different properties, these interactions will be modified together with the structure and its binding attributes will be affected (Alberts B, Johnson A, Lewis J, 2002).

PAML codeml models estimate the impact of the non-synonymous mutations as equal throughout the protein structure. It is important then to differentiate which mutations are affecting the key regions in the protein, resulting into a need of a visual analysis of the location of these mutations in the protein. A python script that reads several files, including the output file from M8 codeml model, and combines them to create a visualization in PyMOL and assess the significance of the selective pressure on the protein domains has been consequently created and presented in this article.

## ALGORITHM DESIGN

The main algorithm behind the script finds the correspondent positions among the gene and the PDB sequences and works in the following manners:

1. Creates 2 lists, list A containing the positions in the alignment (local or global alignment in biopython, 2009) where both sequences don’t have a gap, and list B, which has the length of the gene sequence, filled with ‘nan’ values.
2. Counts the amount of gaps (-, which include not aligned residues) between each segment, bounded by the *i* and *i +1* positions contained in list A, in the aligned sequences (this step is performed in both aligned sequences (aligned chain A and aligned chain B)). This step outputs 2 lists (C and D) with the reciprocal correspondence of the positions where there is not gaps in the alignment of chain A and B.
3. Lists C and D are used to fill in List B taking into account all the indexing issues, PDB (PDB parser, Hamelryck, 2003) and python work in index 1 and 0 respectively, and correspondence. In this step is very easy to substitute the correspondent positions of the gene in the PDB by the actual residues IDs numbers in the PDB structure, which follow their own numbering settings.

## INPUT DATA

The python script requires several input files and counts with several arguments.

## PERSONALIZATION OF PYMOL FEATURES

In order to change and modify the visual aspects of the proteins the script can be easily modified in the PyMOL () function, this are some of the more distinctive features:

### Graphical User Interface (GUI)

The previously described python script has been also embedded into a GUI to facilitate the introduction of input data. The arguments are exactly the same, and the script can be modified internally in the same manners. There are two versions of the script, one of them allows to generate small modifications and remembering the previous file paths given by the user (LYS_PyMOL_GUI_with_temporary_memory.py) due to a loop configuration. The other script (LYS_PyMOL_GUI.py) requires input information every time it is run. LYS_GUI.py only generates the position correlation dataframe, without calling Pymol.

## OUTPUTS

The program will output a 3 column data-frame of the corresponding positions among the gene and the residues IDs in the PDB file. The gene’s positions are organized in a range starting at index 1 and finishing at the length of the sequence (aligned or not). The third column classifies the position according to its belonging to a functional domain (‘Domain’), a selected site(‘Selected’), both (‘Selected_and_Domain’) or none of the previous ones (‘Not’).

This data-frame is given to PyMOL to generate a visualization on the protein from the PDB file information, where the amino acids are colored according to their assigned label.

### Python Script installation requirements

PyMOL installation for the scripts that call PyMOL (Link_Your_Sites_in_PyMOL_Updated.py and LYS_PyMOL_GUI.py), both in python and external (GUI) environments. All the scripts require the python 3 version packages for data manipulation pandas, biopython and numpy. Finally, tkinter for scripts that provide a GUI (LYS_GUI.py and LYS_PyMOL_GUI.py). The scripts are found under https://github.com/artistworking/Master Thesis Scripts

### List of versions of scripts

- **Link_Your_Sites_python_3_Updated.py:** Generates only the text file with the dataframe stating the correspondence of the gene residues positions to the PDB residues positions (according to the PDB special numeration), as seen in Table 3.
- **LYS_GUI.py:** Same as Link_Your_Sites_python_3_Updated.py but with the GUI version.
- **PyMOL_and_LYS_input_dataframe.py:** Simpler script that combines an external dataframe with the same format as the one produced by Link_Your_Sites_python_3_Updated.py (Table 3) and uses it to generate the visualization in PyMOL (Figure 3).
- **Link_Your_Sites_in_PyMOL_Updated.py:** Creates the dataframe in Table 3 and conveys it to PyMOL to generate the visualization in Figure 3.
- **LYS_PyMOL_GUI_Final.py:** The most sophisticated script that does the same as Link_Your_Sites_in_PyMOL_Updated.py but with a GUI. See the tutorial at https://www.youtube.com/watch?v=v3MawxHLGSo

**Figure 1:**
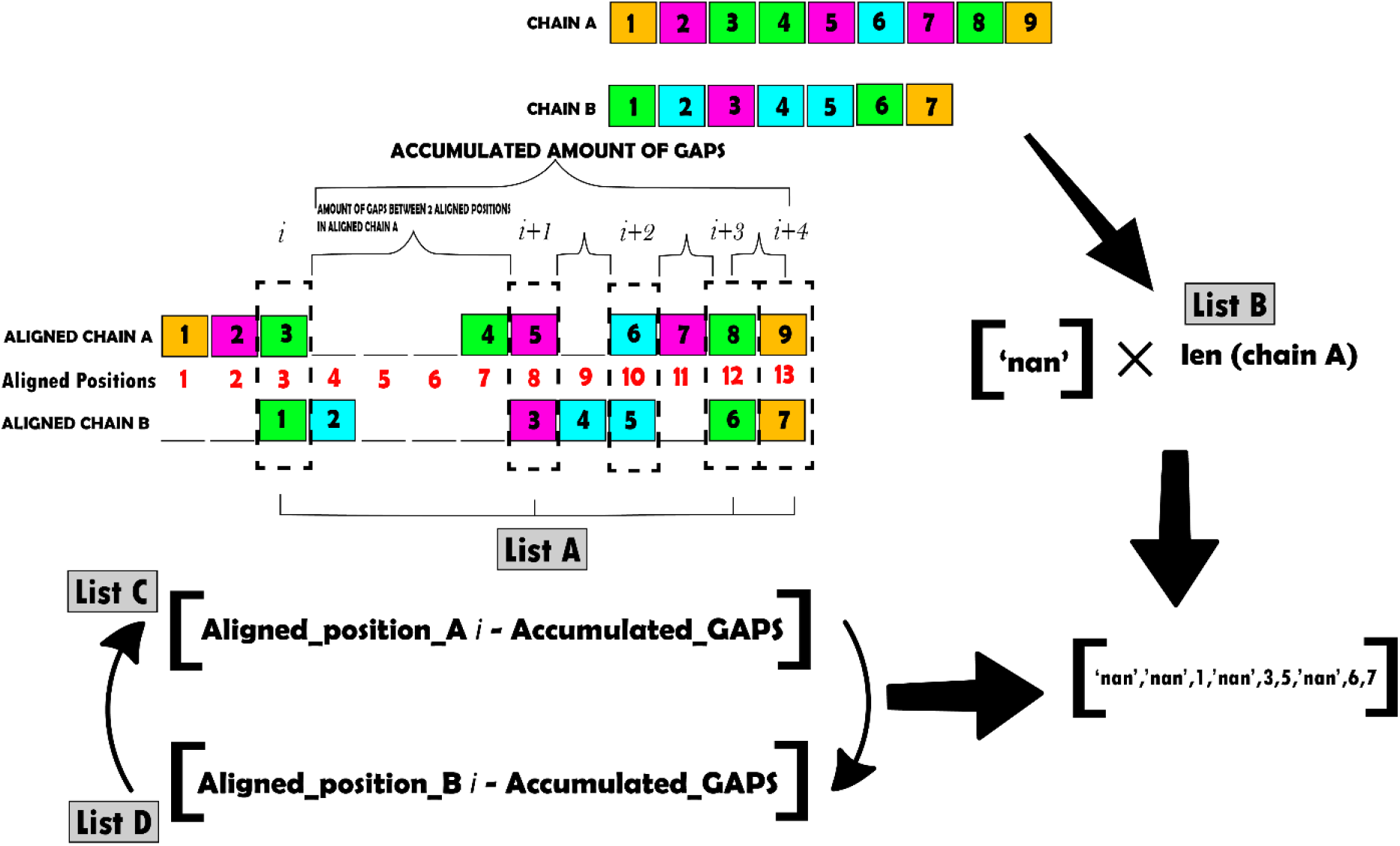
Graphical explanation of the algorithm that matches the coordinates of 2 sequences by using their unaligned and aligned versions (local or global alignment in biopython, 2009). The numbers indicate the residues positions in the chain/sequence.

**Figure 2:**
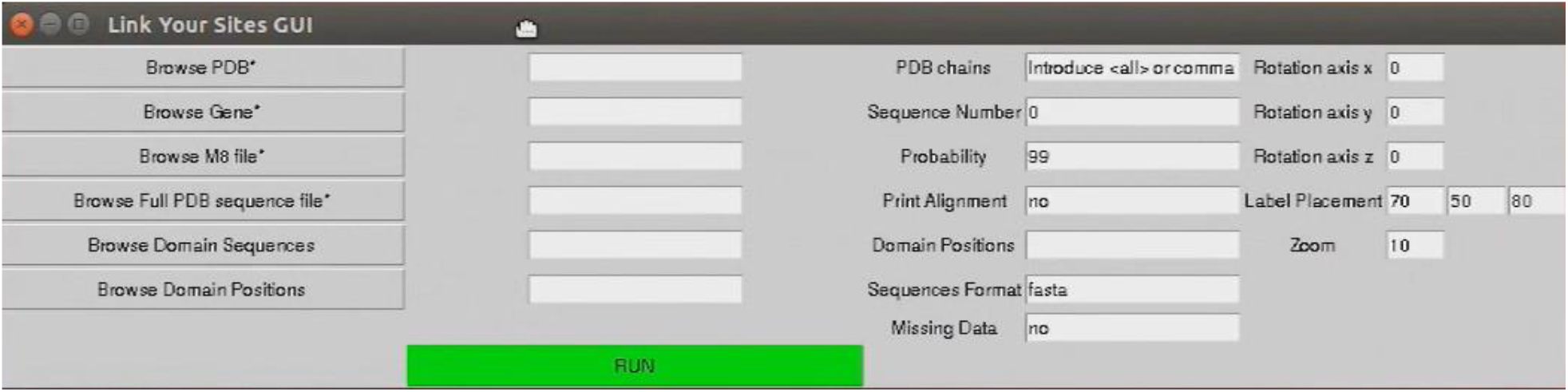
LYS.py interface. The compulsory files for the GUI to work are marked with a *. Check an introductory tutorial at https://www.youtube.com/watch?v=v3MawxHLGSo

**Figure 3.**
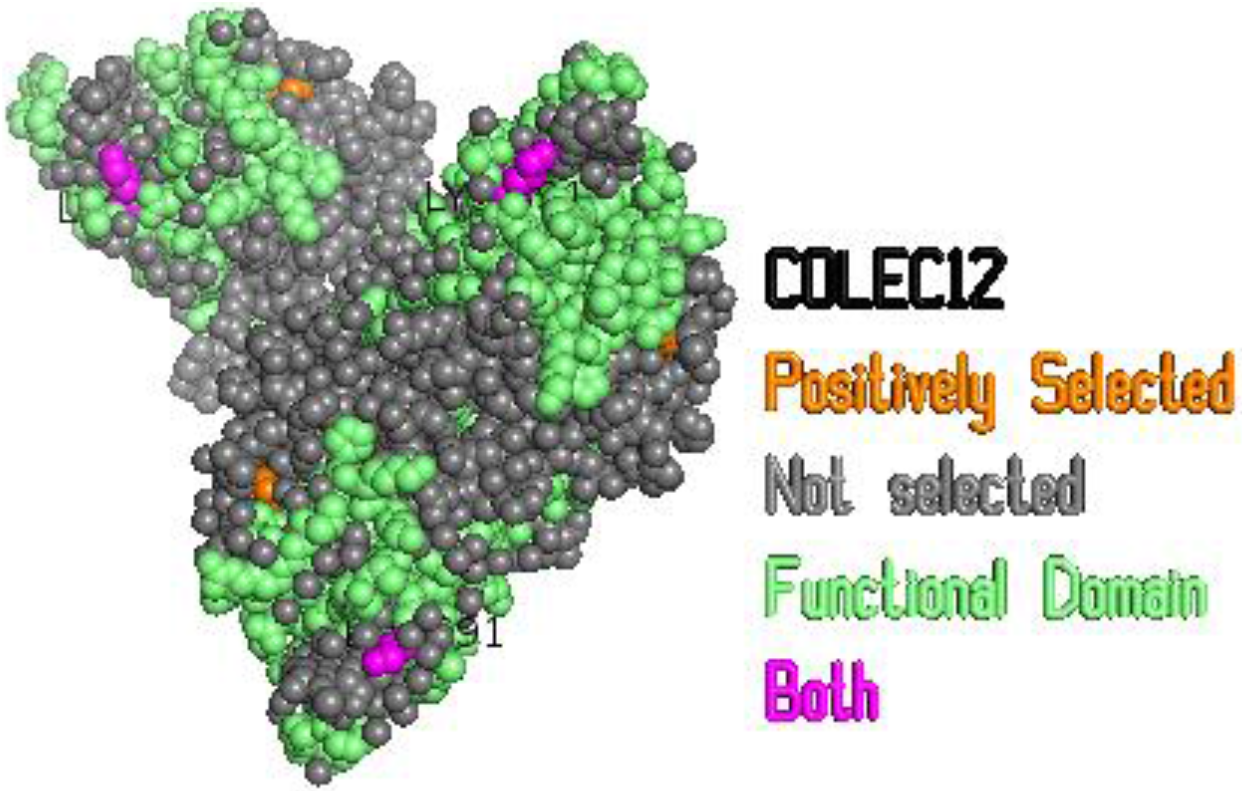
LYS.py visual output example of protein coloured according to its evolutionary positively selected amino acid residues and domain positions.

**Table 1.**
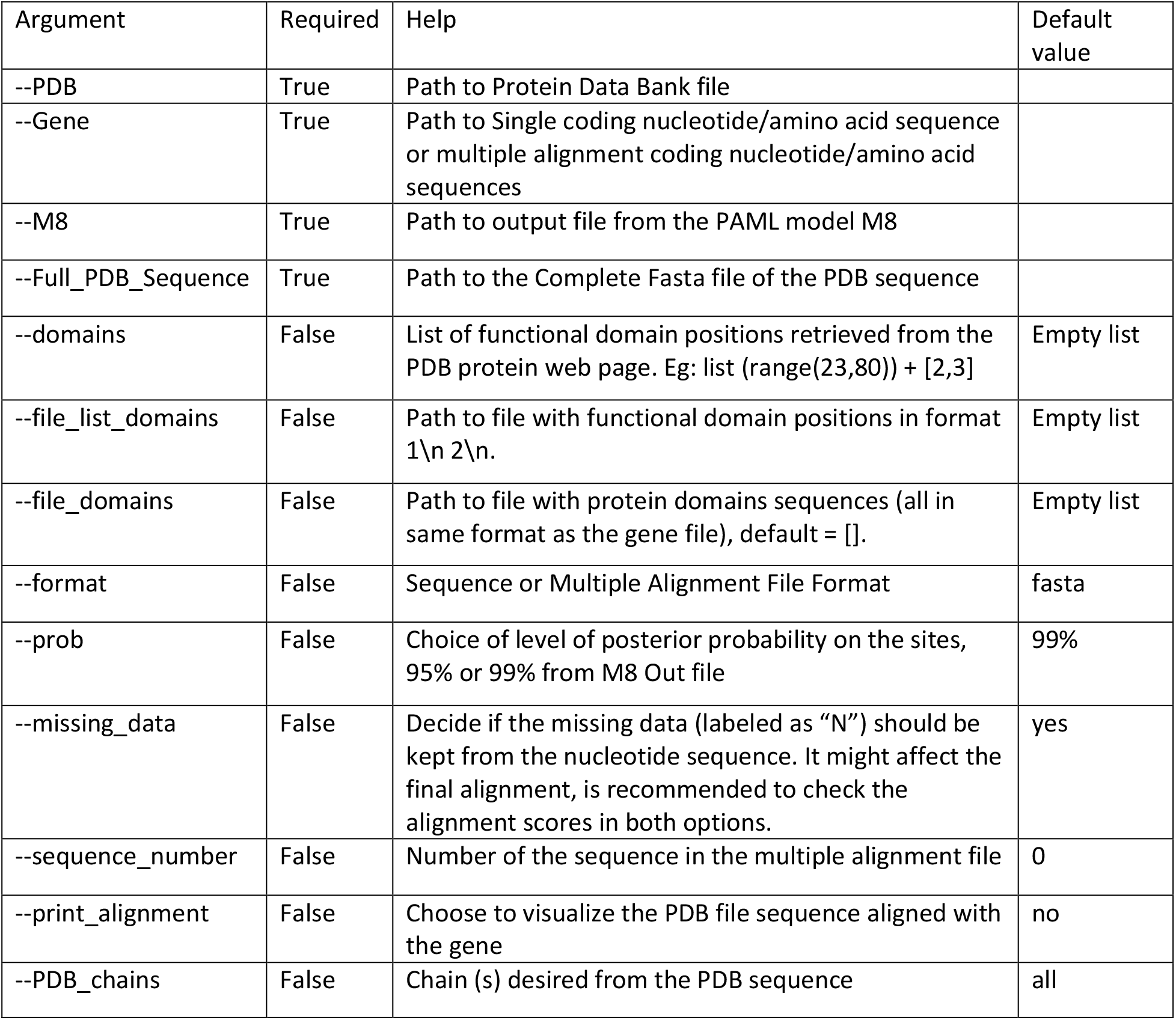
LYS.py script list of arguments.

| Argument | Required | Help | Default value |
| --- | --- | --- | --- |
| --PDB | True | Path to Protein Data Bank file |  |
| --Gene | True | Path to Single coding nucleotide/amino acid sequence or multiple alignment coding nucleotide/amino acid sequences |  |
| --M8 | True | Path to output file from the PAML model M8 |  |
| --Full_PDB_Sequence | True | Path to the Complete Fasta file of the PDB sequence |  |
| --domains | False | List of functional domain positions retrieved from the PDB protein web page. Eg: list (range(23,80)) + [2,3] | Empty list |
| --file_list_domains | False | Path to file with functional domain positions in format 1\n 2\n. | Empty list |
| --file_domains | False | Path to file with protein domains sequences (all in same format as the gene file), default = []. | Empty list |
| --format | False | Sequence or Multiple Alignment File Format | fasta |
| --prob | False | Choice of level of posterior probability on the sites, 95% or 99% from M8 Out file | 99% |
| --missing_data | False | Decide if the missing data (labeled as "N") should be kept from the nucleotide sequence. It might affect the final alignment, is recommended to check the alignment scores in both options. | yes |
| --sequence_number | False | Number of the sequence in the multiple alignment file | 0 |
| --print_alignment | False | Choose to visualize the PDB file sequence aligned with the gene | no |
| --PDB_chains | False | Chain (s) desired from the PDB sequence | all |

**Table 2:** LYS.py customizable features in the script.

| Settings | Options |
| --- | --- |
| Background, Residues and Font Colours | Choose colours from the palette:<br>https://pymolwiki.org/index.php/Color_Values. |
| Residues Shapes | Choose from:<br>https://pymolwiki.org/index.php/Show |
| Select and Remove chains | Chose if any of the chains should be removed in the visualization. |
| Protein Rotation | Rotate the protein accordingly in the [ x, y,z ] axis. |
| Legend : Font Size and Placement | Change the values of the axes, cyl_text and cmd.set |
| Carbon-alpha residues labelling | Activate cmd.label accordingly: Designed to highlight only alpha carbons of selected sites |

**Table 3:** LYS.py table output of correspondence among the coordinates/residues of the studied sequences

| Gene_Position | PDB_Position | Label |
| --- | --- | --- |
| 1 | Nan | Not |
| 2 | -1 | Domain |
| 3 | 0 | Selected |
| 4 | 432 | Selected_and_Domain |

